# Multi-modal microvascular cerebral blood flow velocity mapping with 14T single-vessel MRI and optical microscopy in the mouse brain

**DOI:** 10.1101/2023.11.07.566001

**Authors:** D. R. Miller, X. A. Zhou, S. Liang, Z. Xie, B. Fu, P. Shin, Q. Pian, Y. Jiang, A. Liu, J. Tang, A. Devor, X. Yu, S. Sakadzic

## Abstract

In this study, we imaged the same penetrating cortical vessels in a mouse using ultrahigh field single-vessel MRI at 14 T and high-resolution optical microscopy imaging. The optical imaging was performed through a chronic sealed cranial window, while the single-vessel MRI was facilitated by a custom-designed, chronically implanted radiofrequency coil surrounding the window. The MRI and optical imaging were performed sequentially focused on the same penetrating cortical arterioles and surfacing venules within the whisker barrel cortex. With MRI, we obtained high-resolution multi-gradient echo (MGE) images and single-vessel phase contrast (PC) velocity maps. With optical imaging, we acquired microvascular angiograms using 2-Photon Microscopy (2PM) and Optical Coherence Tomography (OCT) and measured the blood flow velocity using Dynamic Light Scattering OCT (DLS-OCT). The MGE images, PC-based MRI velocity maps, OCT angiograms, and DLS-OCT velocity maps were coregistered with the 2PM microvascular angiograms. Using these tools, we cross-validated blood flow velocity in the penetrating cortical arterioles and surfacing venules measured by single-vessel MRI and OCT at rest. Our novel method demonstrates the possibility of combining ultrahigh field single-vessel MRI and high-resolution optical methods (e.g., 2PM and OCT) for studying brain structure and function with single microvessel precision.

## 1. Introduction

The brain’s vascular network plays a pivotal role in maintaining its intricate functions, providing oxygen and nutrients to the brain tissue, and enabling waste removal. Understanding the dynamics of blood flow at the single-vessel level, such as within penetrating arterioles and venules, is crucial for understanding the microvascular organization and neurovascular mechanisms in health and disease. However, gaining insights into vascular dynamics at the single-vessel scale has remained a challenge due to vessel size and the complexity of vessel interactions.

In animals, microscopic optical imaging has generated a body of detailed information about microvascular structure and dynamics at the level of single vessels [1-4]. In humans, magnetic resonance imaging (MRI) has been widely used for angiography, measurements of blood flow (i.e., using arterial spin labeling) and inference of neuronal activity through hemodynamics (i.e., BOLD fMRI). Previously, we have shown that structural and fMRI can be performed in mice implanted with chronic cranial optical windows opening the door for longitudinal multimodal optical/MR “preclinical” imaging studies with translational potential [5]. Others have combined fMRI with simultaneous mesoscopic optical imaging [6, 7]. A proof-of-concept also has been achieved for simultaneous 2PM and fMRI imaging by employing free propagation of the 2PM excitation and fluorescence emission light between the control room and the MRI scanner [8].

Previously, we developed the single-vessel fMRI method with spatial resolution of 20-70 μm in rodent brain [9]. This method allows mapping of the cerebral blood flow, volume, and oxygenation changes in single penetrating cortical arterioles and surfacing venules [9-12], bridging preclinical fMRI to the scale of optical microscopy. Critically, we have improved the signal-to-noise ratio (SNR) of fMRI signals by developing chronically implanted radiofrequency (RF) coils [12]. However, *in vivo* validation of single-vessel fMRI measurements remains an outstanding task.

In contrast with fMRI, optical microscopy tools offer direct, model-free measurements of the vascular parameters of interest. In particular, Optical Coherence Tomography (OCT) enables direct measurement of blood flow velocity in the cortical microvasculature [2, 4], and Two-Photon Microscopy (2PM) provides high-resolution three-dimensional microvascular morphology [1, 3]. Together, OCT and 2PM offer complementary information needed to validate fMRI at the single-vessel level. In practice, 2PM and OCT imaging of a mouse cortex are typically performed through a sealed cranial window. Therefore, using 2PM and OCT for validation of fMRI single-vessel measurements in the same mouse is a challenging task that requires combining the window assembly with the implanted RF coil. An additional challenge is accurate coregistration of 3-dimensional (3D) data sets obtained across imaging modalities.

Here, we solve these challenges enabling validation of single-vessel MRI velocity measurements in the same mouse using Dynamic Light Scattering (DLS) OCT. We combined an implantable RF coil with a 3-mm diameter chronic cranial window enabling chronic high-resolution imaging of the brain vasculature with 2PM, OCT, and single-vessel MRI. We used anatomical imaging of vessels obtained by each modality to coregister the multimodal data. We coregistered the maps of arterioles and venules (“A-V maps”) based on multi-gradient echo (MGE) MRI with 2PM vascular angiograms using pial vessels visible in both data sets. Similarly, we coregistered OCT angiograms of cortical vasculature with the same 2PM vascular angiograms that were coregistered with the MRI A-V maps. Immediately after acquiring MRI A-V maps, we also obtained phase contrast (PC) MRI flow maps to quantify the blood flow velocity of penetrating arterioles and surfacing venules. These MRI flow maps are inherently spatially coregistered with the MRI A-V maps and, therefore, coregistered with the 2PM vascular angiograms. We also performed DLS-OCT immediately after obtaining OCT angiograms to quantify the blood flow velocity in penetrating arterioles and surfacing venules. The DLS-OCT blood flow velocity maps are inherently spatially registered with the OCT angiograms, and thus also coregistered with the 2PM vascular angiograms. Finally, we quantified the blood flow velocity in the same penetrating arterioles and surfacing venules with PC MRI and DLS-OCT. Our study proves the feasibility of validating high-resolution MRI-based vessel velocity measurements with optical microscopy and demonstrates a potential to use fMRI for extending quantitative single-vessel measurements beyond the depth penetration limit of optical methods.

## 2. Materials and Methods

### 2.1A Animal Procedures

All animal surgical and experimental procedures were conducted following the Guide for the Care and Use of Laboratory Animals and approved by the Massachusetts General Hospital Subcommittee on Research Animal Care. We used healthy 6–12-month-old female C57BL/6 mice (The Jackson Laboratory, Maine, United States). The mice underwent craniotomy surgery to install an MRI coil surrounding a chronic sealed cranial window for optical access to the whisker barrel cortex on the left hemisphere. A 3-mm-diameter circular area was exposed with center coordinates of 2 mm posterior to bregma and 3 mm lateral from the midline. The area was then sealed with a glass window implant using instant adhesives (Loctite 4014 and 401; Henkel); dental acrylic was not used to avoid interference with the MRI signal quality. After installation of the cranial window, a custom-built MRI coil was positioned over the cortex and tested for tuning and matching signal with a network analyzer. The MRI coil was positioned to maintain optical access to the barrel cortex. The coil ring was lifted ∼0.5 mm above the surface of the skull and held in place while a thin layer of cyanoacrylate adhesive (Loctite 404, Henkel) was applied to connect the skull with the coil. Once the adhesive had dried (∼8 mins), 2-part dental cement was applied to cover the coil and exposed bone, paying special attention to firmly secure the base of the coil. Finally, the edges of the skin were attached using the cyanoacrylate adhesive (Loctite 454, Henkel). After the dental cement had fully hardened (∼10 mins), the mouse was released from the stereotaxic frame and allowed three weeks to recover fully from surgery. **Fig. 1A** shows an example of the cranial window after installing the MRI coil. The MRI coils are sufficiently small to not obstruct the access of the high-magnification multiphoton objectives (**Fig. 1B**). During optical and MRI imaging sessions, mice were anesthetized with the mixture of medical air (1.1 L/min), oxygen (0.3 L/min), and isoflurane (1.5%). A heating pad was used to ensure the mouse core body temperature was maintained at a range between 36–37°C throughout anesthesia.

**Fig. 1.**
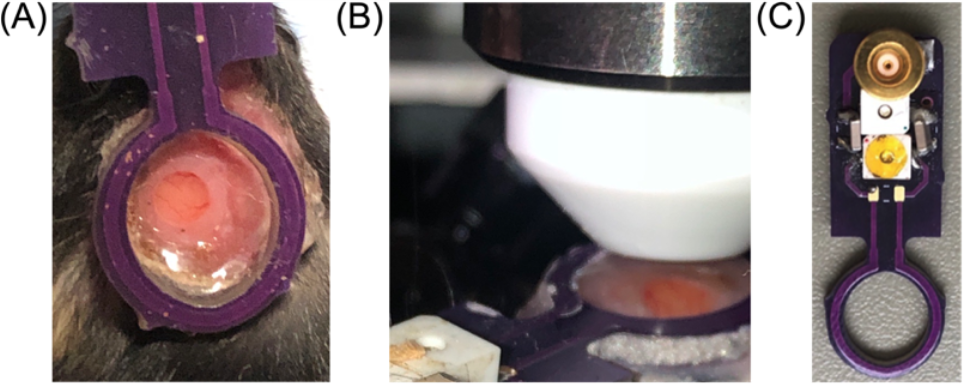
**(A)** Example of cranial window of mouse after MRI coil installation surgery. **(B)** Demonstration of high-magnification multiphoton objective positioned over the mouse cortical surface. **(C)** MRI transceiver coil.

### 2.1B MRI coils

The MRI transceiver coil was designed and built as seen in **Fig. 1C**. The coils were built to optimize impedance matching on a mouse head, and the surgical attachment serves to minimize the air-tissue interface. Each coil was built to weigh ∼2.5 g to minimize the effect on animal motion and facilitate recovery and neck strengthening for each mouse. The design keeps the B1 direction orthogonal to the B0. The single loop allows full brain coverage at sufficient depths for various study paradigms.

### 2.2 Imaging Methods

For each mouse (*n*=2), we performed separate imaging sessions for 1) MRI, 2) 2PM, and 3) OCT. At least one day of recovery was allowed between different imaging sessions. The same anesthesia regime, as described in Sec. 2.1, Animal Procedures, was applied during all imaging sessions. During imaging, mice were head-restrained via a custom-built holder for the MRI coil.

### 2.2A Magnetic Resonance Imaging

MRI was performed by using a 14T/6 cm horizontal-bore magnet (Magnex Scientific) interfaced through the Bruker Avance III (Bruker). The scanner has a 12 cm magnet gradient set with a strength of 100 G/cm, and a 150 μs rise time (Resonance Research Inc.). Homemade surface transceiver RF coils with an internal diameter of 7.5 mm were used for MRI image acquisition.

#### Single-vessel MGE Imaging

For the detection of the individual arterioles and venules in the cortex, a 2D multi gradient echo (MGE) sequence was applied with the following parameters: TR: 100 ms; TE: 2.5, 5.5, 11.5, 14.5, 17.5, 17.5, 20.5, 23.5 ms; flip angle: 58°; matrix: 200 × 200; in-plane resolution: 40 × 40 μm^2^; slice thickness: 200 μm. The MGE images were averaged from the 1st echo to the 5th/6th echo to get the A–V map. We obtained MGE MRI images for axial depths from 0-200, 200-400, 400-600, and 600-800 μm below the surface of the mouse cortex. The images were acquired such that the axial axis was perpendicular to the sealed cranial window.

#### Flowmap

For the measurement of axial cerebral blood flow (CBF) velocity in the same field of view (FOV) of the MGE, PC MRI was applied with the following parameters: TR: 50 ms; TE: 4.4 ms; flip angle: 58°; matrix: 160 × 160; in-plane resolution: 50 × 50 μm^2^; slice thickness: 300 μm. Single-vessel MRI blood flow velocity maps were then acquired at axial depths from 50-350 and 350-650 μm below the cortical surface. The blood flow velocity images have 50 μm lateral resolution, 300 μm axial resolution, and a maximum measurable absolute blood flow velocity of 15 mm/s.

### 2.2B Two-Photon Microscopy (2PM)

For *in vivo* 2PM, we labeled the blood plasma with dextran-conjugated Alexa-680 (70 kDa). We imaged microvasculature over the entire 3-mm-wide cranial window with a low-magnification 4X objective (XLFLUOR4X, Olympus) to an axial depth of 800 μm below the cortical surface using 2 μm axial steps; each axial slice is 512×512 pixels equating to 7 μm steps per XY pixel. We used mode-locked, pulsed excitation laser light at λ=1,280 nm (Chameleon Discovery NX, Coherent, 100 fs pulse width, 80 MHz pulse repetition rate). Laser power was controlled by an electro-optic modulator (EOM) (350-105-02; Conoptics Inc.). Transverse scanning was performed by a two-axis 7 mm galvanometer scanner (6210 H; Cambridge Technology). The laser beam was relayed to the back focal plane of the objective through the combination of a scan lens (*f* = 51 mm, 2x AC508-100-B, Thorlabs) and a tube lens (*f* = 180 mm, Olympus). The axial movement of the objective was controlled by two translation stages (ZFM2020 and PFM450E; Thorlabs). Fluorescence emission signal from Alexa-680 was collected by a photomultiplier tube (PMT) (H10770PA-50; Hamamatsu) after passing through a dichroic mirror (FF875-Di01-38.1×51; Semrock Inc) and two fluorescent filters (FF01-890/SP-50 and FF01-709/167-25; Semrock Inc).

### 2.2C Dynamic Light Scattering Optical Coherence Tomography (DLS-OCT)

We utilized a low-coherence superluminescent diode light source with a center wavelength of λ=1,325 nm (LS2000B; Thorlabs) and a bandwidth exceeding 170 nm. Our system incorporated a line scan camera with a 46-kHz acquisition rate (SU1024-LDH2; Sensors Unlimited). The setup enabled a maximum measurable absolute blood flow velocity of ∼10 mm/s, without phase wrapping. The minimum detectable absolute velocity was 0.7 mm/s, influenced by the system’s phase noise measured at ±0.2 radians. For our imaging experiments, we used a 10X objective lens (M Plan Apo NIR for Bright Field Observation 378-823-5; Mitutoyo), achieving a transverse resolution of 3.5 μm. The system offers an axial resolution of 2.7 μm in cortical tissue for a depth range of up to 1 mm. The incident optical power on the sample was 2 mW.

To calculate axial blood flow velocity with DLS-OCT, we used the first-order field autocorrelation function analysis-based OCT velocimetry method (g_1_V_z_) using a similar protocol as reported previously by Lee et al. (2012), Tang et al. (2018), and Guo et al. (2023) [13-15]. The volume size was 3 mm × 3 mm, and each volume was acquired with 600 B-scans, where 600 A-scans were acquired per B-scan and 100 repeats for each lateral position. It took approximately 15 minutes for data acquisition. For comparing axial blood flow velocity with single-vessel fMRI measurements, we selected the maximum axial blood flow velocity value in the 3D DLS-OCT volumes at the centers of the penetrating arterioles and venules. The OCT angiograms were acquired in the same experimental session with the DLS-OCT velocity maps. This ensured that the DLS-OCT velocity maps are inherently spatially registered with the OCT angiograms (**Supplemental Fig. S1)**, which facilitated coregistration of the OCT and 2PM images.

### 2.3 Coregistration of Multimodal Images

To coregister the multimodal imaging data, we first resampled 2PM angiograms in 3D space to reduce effects from curvature of the brain surface and any tilt of the animal during 2PM imaging. The resampled angiograms then served as the coordinate space to which MRI and OCT measurements were coregistered. MRI MGE and MRI single-vessel PC velocity maps were obtained consecutively during the same experiment; thus, the structural information contained in the MRI MGE images is inherently spatially coregistered with the velocity maps (**Fig. 3**).

**Fig. 3.**
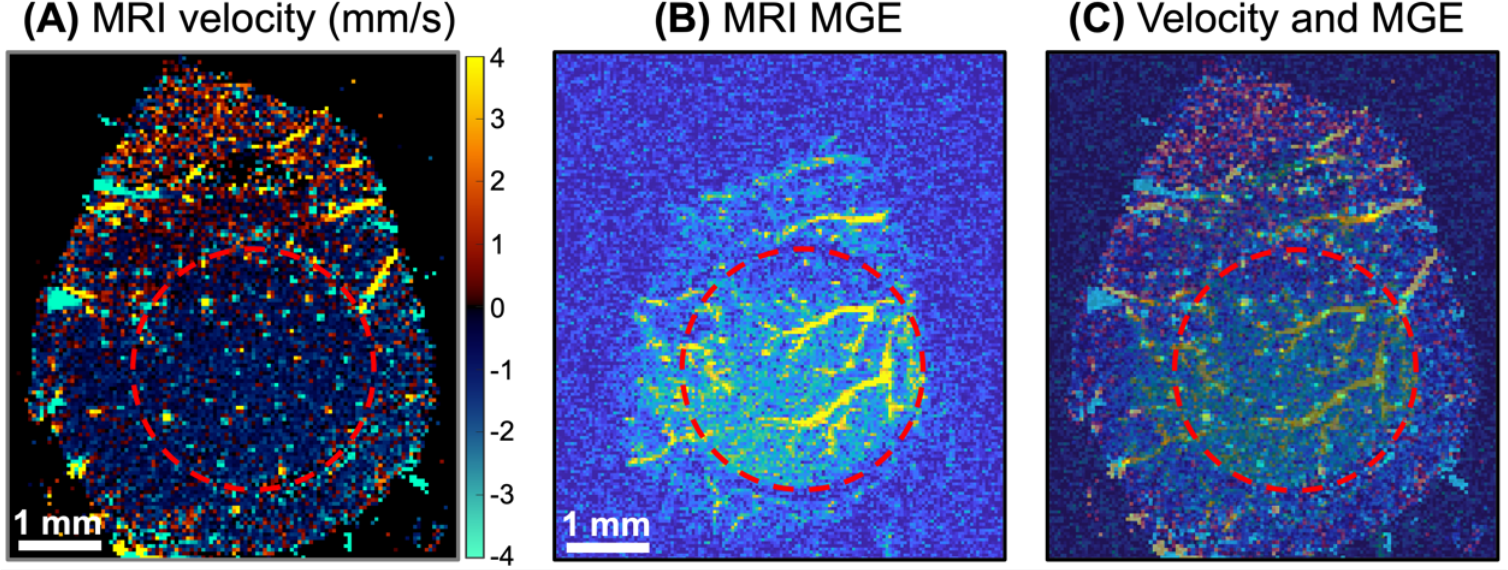
An example of the MRI single-vessel phase contrast velocity map **(A)** and MRI multi-gradient echo (MGE) structural image **(B)**. The velocity map is a single slice from 50-350 μm below the cortical surface and the MGE is a single slice from 0-200 μm below the cortical surface. The MRI velocity map and MGE image are overlaid in panel **(C)**. To aid the viewer, a red dashed circle has been added to each panel outlining the edge of the 3 mm diameter cranial window.

We first coregistered an MRI MGE slice from 0-200 μm below the cortical surface with the resampled 2PM angiogram. Large vessels visible in the maximum intensity projection (MIP) of the 2PM angiogram and the MRI MGE image were used as fiducial markers. Multiple points were selected based on the fiducial markers to calculate an affine transformation to map the MRI MGE slice to the 2PM angiogram. In a similar manner, *en face* projections of the OCT angiogram were coregistered with the MIP of the 2PM angiogram using visible vessels as fiducial markers. An example coregistering a 2PM angiogram, OCT angiogram, and MRI MGE is shown in **Fig. 4**. Using this method, MRI MGE images, MRI velocity maps, OCT angiograms, and DLS-OCT velocity maps were all coregistered with the 2PM angiogram and thus coregistered with each other.

**Fig. 4.**
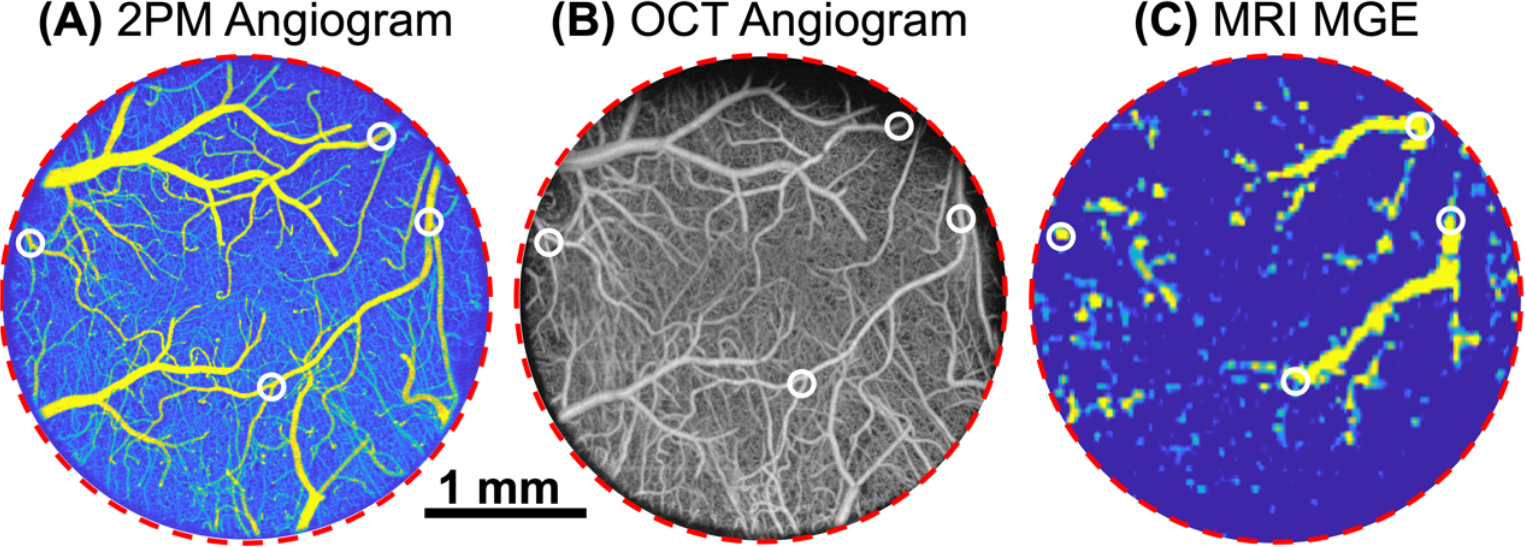
Example of co-registration of **(A)** 2PM angiogram maximum intensity projection from 0-600 μm below the cortical surface, **(B)** OCT angiogram maximum intensity projection from 0-500 μm below the cortical surface and **(C)** MRI MGE maximum intensity projection from 0-200 μm below the cortical surface. Four points identified by white circles are coregistered in each panel. To aid the viewer, a red dashed circle has been added to each panel outlining the edge of the 3 mm diameter cranial window. Scale bar is 1 mm and applies to each panel.

## 3. Results and Discussion

We compared axial blood flow velocities in the penetrating arterioles and surfacing venules in the mouse cortex measured by single-vessel PC MRI and DLS OCT for *n*=2 mice at rest. **Figs. 5** and **6** show an example of coregistered velocity maps of penetrating arterioles and surfacing venules, respectively, for one mouse. **Supplemental Figs. S2** and **S3** show coregistered velocity maps of penetrating arterioles and surfacing venules for the second mouse.

**Fig. 5.**
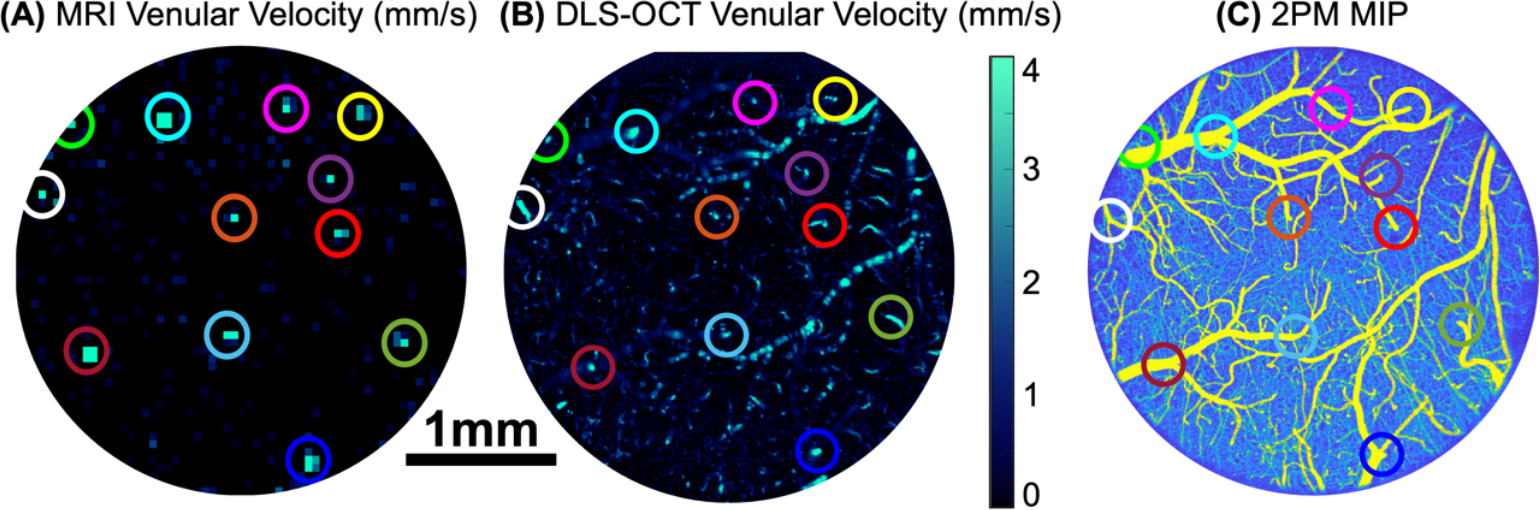
Comparison of venular axial velocity maps from **(A)** PC MRI and **(B)** DLS-OCT. The PC MRI velocity map is a single slice from 50-350 μm below the cortical surface and the DLS-OCT velocity map is a maximum intensity projection (MIP) from 0-500 μm which is why pial vessels are visible. The selected points are also shown in the 2PM microvascular angiogram shown as a MIP from 0-600 μm below the cortical surface **(C)**. The same circle colors in all three panels are used to mark the same surfacing venules. Scale bar is 1 mm and applies to each panel.

**Fig. 6.**
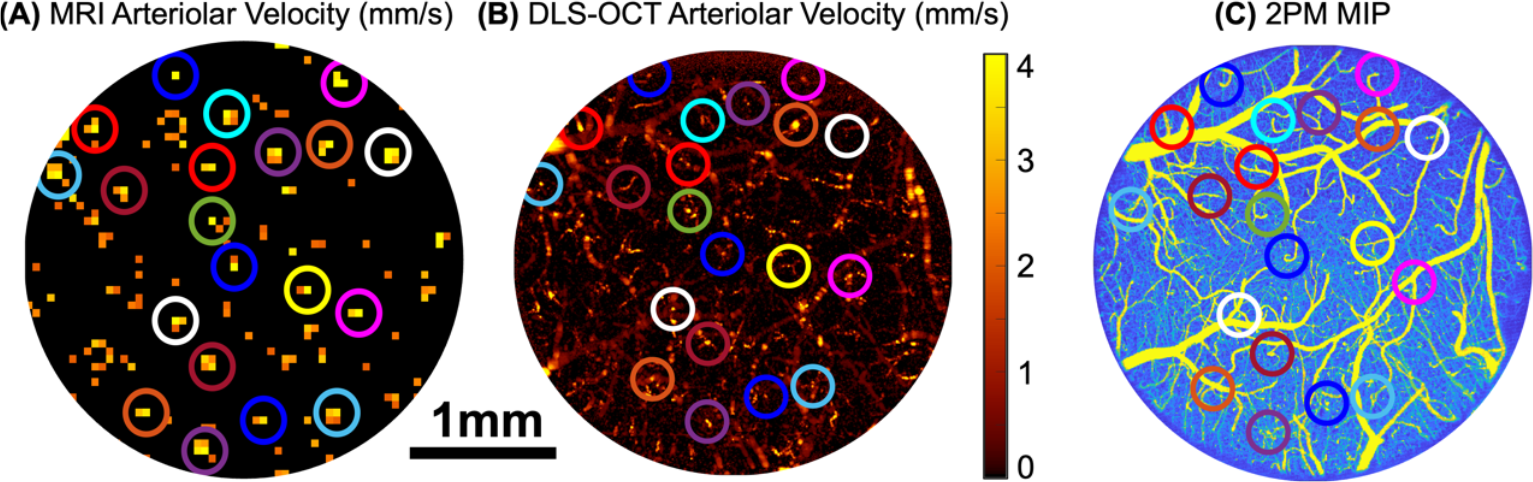
Comparison of arteriolar axial velocity maps from **(A)** PC MRI and **(B)** DLS-OCT. The PC MRI velocity map is a single slice from 50-350 μm below the cortical surface and the DLS-OCT velocity map is a maximum intensity projection (MIP) from 0-500 μm which is why pial vessels are visible. The selected points are also shown in the 2PM microvascular angiogram shown as a MIP from 0-600 μm below the cortical surface **(C)**. The same circle colors in all three panels are used to mark the same diving arterioles. Scale bar is 1 mm and applies to each panel.

Following the coregistration procedures, the same penetrating vessels visible in the MRI velocity maps and the DLS-OCT velocity maps were identified. For both PC MRI and DLS-OCT velocity maps, the pixel with the highest velocity within the vessel was used. The axial velocities of each selected vessel from the MRI and DLS-OCT velocity maps were tabulated and the vessel diameter was estimated based on the 2PM angiogram. We measured the axial velocities in a total of 66 vessels from *n*=2 mice, of which 35 are arterioles and 31 are venules. **Fig. 7** shows a box plot of the axial velocities for all selected arterioles (**Fig. 7A**) and venules (**Fig. 7B**). The median arteriolar axial velocities measured by DLS-OCT and MRI were 5.6 mm/s and 3.8 mm/s, respectively. The median venular axial velocities measured by DLS-OCT and MRI were 7.1 mm/s and 4.1 mm/s, respectively. Lower median arteriolar and venular axial velocities observed in MRI measurements may be a result of the different spatial resolutions of MRI and DLS-OCT and small diameters of the penetrating cortical vessels in mice. The majority of the measured penetrating vessels (70%) had diameters smaller than 50 μm, which is the transverse resolution of PC MRI velocity maps in our experiments. Therefore, the pixel with the largest velocity value in the MRI data in most cases represented the average over the distribution of velocity values across the cross-section of the penetrating vessel. Due to the blunted parabolic profile of the red blood cell velocities across the vessel cross-section [14, 16], the average velocity may be significantly lower than the maximal velocity along the axis of the vessel. On the other side, the maximum blood flow velocity estimated by DLS-OCT in penetrating vessels is likely much closer to the true maximal velocity due to the high transverse resolution of the DLS-OCT measurements (e.g., 3.5 μm), which is much smaller than the diameter of the penetrating vessels.

**Fig. 7.**
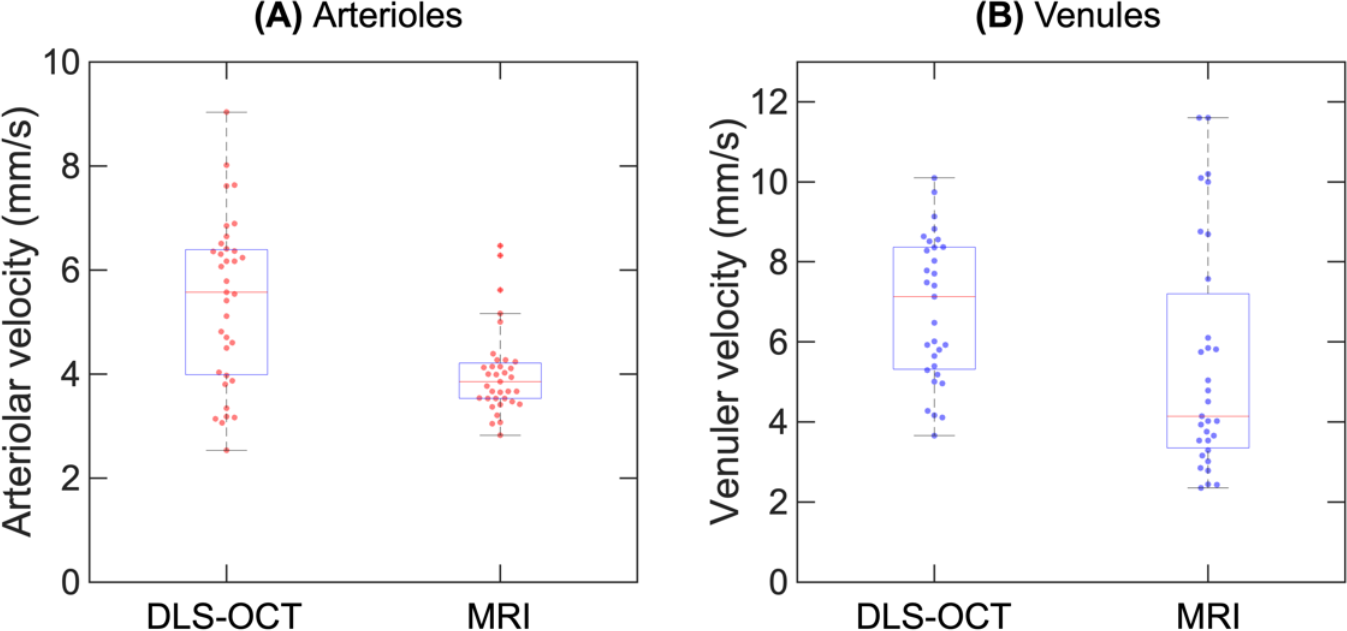
Comparison of the **(A)** arteriolar axial blood flow velocities and **(B)** venular axial blood flow velocities measured by DLS-OCT and PC MRI. In the box plots, arteriolar and venular velocity measurements are shown using red and blue symbols, respectively.

The discrepancy in spatial resolutions of single vessel PC MRI and DLS-OCT imposed a limitation in this study to the exact comparison of maximal blood flow velocities measured by these two modalities. However, the availability of the high-resolution microvascular angiograms obtained by 2PM in combination with the known blunted parabolic profiles of the blood flow inside the vessels, will enable us to closely analyze the single-vessel MRI measurements of blood flow velocities in microvessels with various diameters. Another limitation of the current work is that measurements were performed in anesthetized animals. Potential variations in the anesthesia level during different imaging sessions may have affected both absolute blood flow velocity and vascular diameters. However, the developed methodology of animal preparation and data processing should be directly applicable to single-vessel MRI and optical imaging in awake mice. Moreover, future studies could easily incorporate single-vessel fMRI (i.e., multiple time points with the same procedures described herein), single-vessel fMRI BOLD and blood volume imaging, and high-resolution optical imaging of the intravascular oxygenation.

## 4. Conclusions

We developed a novel methodology to compare axial blood flow velocities in the same penetrating arterioles and surfacing venules based on single-vessel PC MRI and DLS-OCT measurements. To this end, we optimized the design of the MRI coil permanently fixed to the mouse head in combination with the sealed cranial window to enable chronic imaging of the same cortical region by both single-vessel PC MRI and high-resolution optical microscopy (e.g., OCT and 2PM). We further developed the algorithms to accurately coregister MRI and DLS-OCT measurements with the help of 2PM angiograms and identify the same penetrating arterioles and surfacing venules in all data sets. We used these tools to identify 66 penetrating vessels in *n*=2 mice and perform a first direct comparison of the axial blood flow velocity measurements obtained by single-vessel PC MRI and DLS-OCT. Future studies will utilize the developed tools to better understand blood flow velocity measurements by single-vessel fMRI in vascular segments with different vessel diameters, and to validate single-vessel fMRI measurements of the blood volume and BOLD signal in both anesthetized and awake animals.

## Acknowledgements

We thank Dr. Kivilcim Kiliç for assistance with resampling 2PM angiograms.

## Supplementary Information

**Supplemental Fig. S1** shows the spatial overlap of a DLS-OCT velocity map and OCT angiogram.

**Fig. S1.**
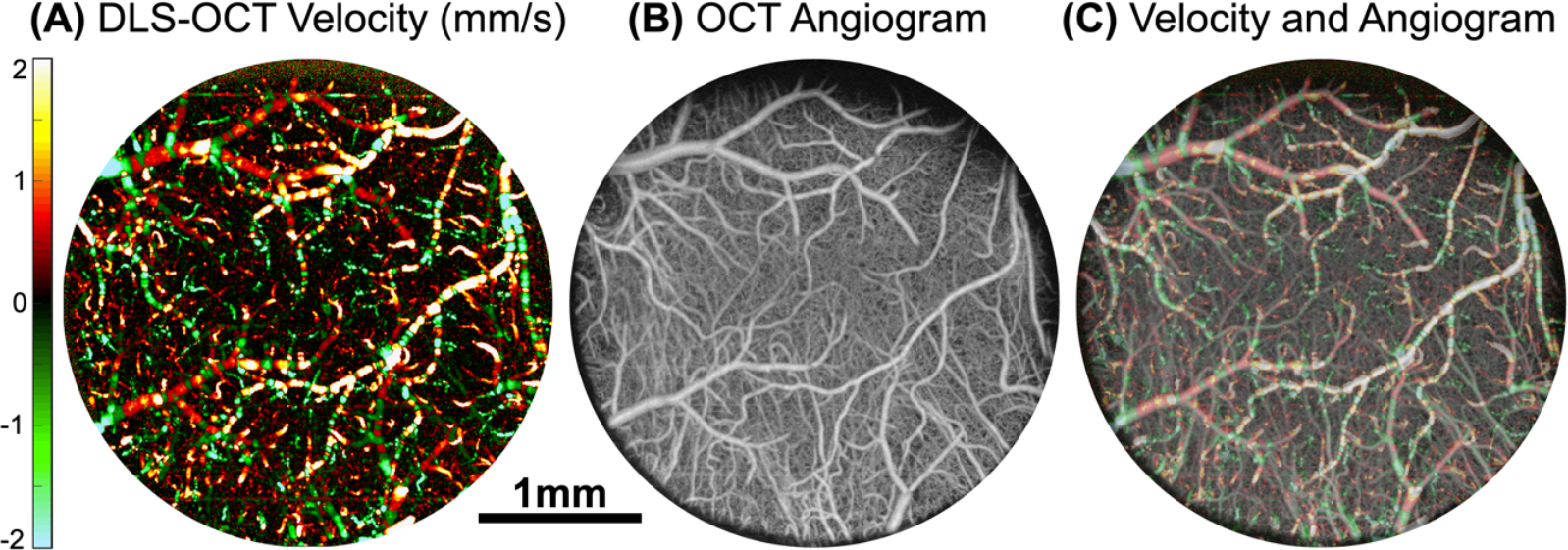
Demonstration of the DLS-OCT velocity map **(A)** and OCT angiogram **(B)**. Both the DLS-OCT velocity map and angiogram are maximum intensity projects from 0-500 μm below the cortical surface. The DLS-OCT velocity map and OCT angiogram are overlaid in panel **(C)**.

**Supplemental Figs. S2** and **Fig. S3** show a comparison of the surfacing venules and penetrating arterioles, respectively, in the second mouse.

**Fig. S2.**
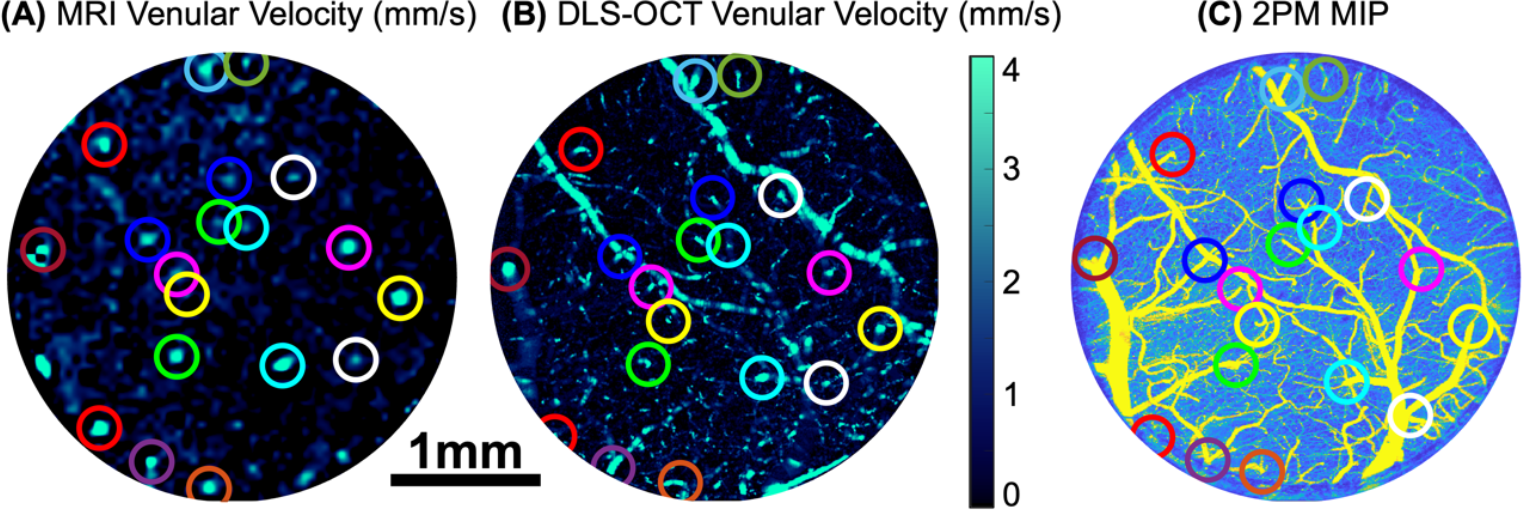
Comparison of venular axial velocity maps from **(A)** PC MRI and **(B)** DLS-OCT for the second mouse. The PC MRI velocity map is a single slice from 50-350 μm below the cortical surface and the DLS-OCT velocity map is a maximum intensity projection (MIP) from 0-500 μm which is why pial vessels are visible. The selected points are also shown in the 2PM microvascular angiogram shown as a MIP from 0-600 μm below the cortical surface **(C)**. The same circle colors in all three panels are used to mark the same surfacing venules. Scale bar is 1 mm and applies to each panel.

**Fig. S3.**
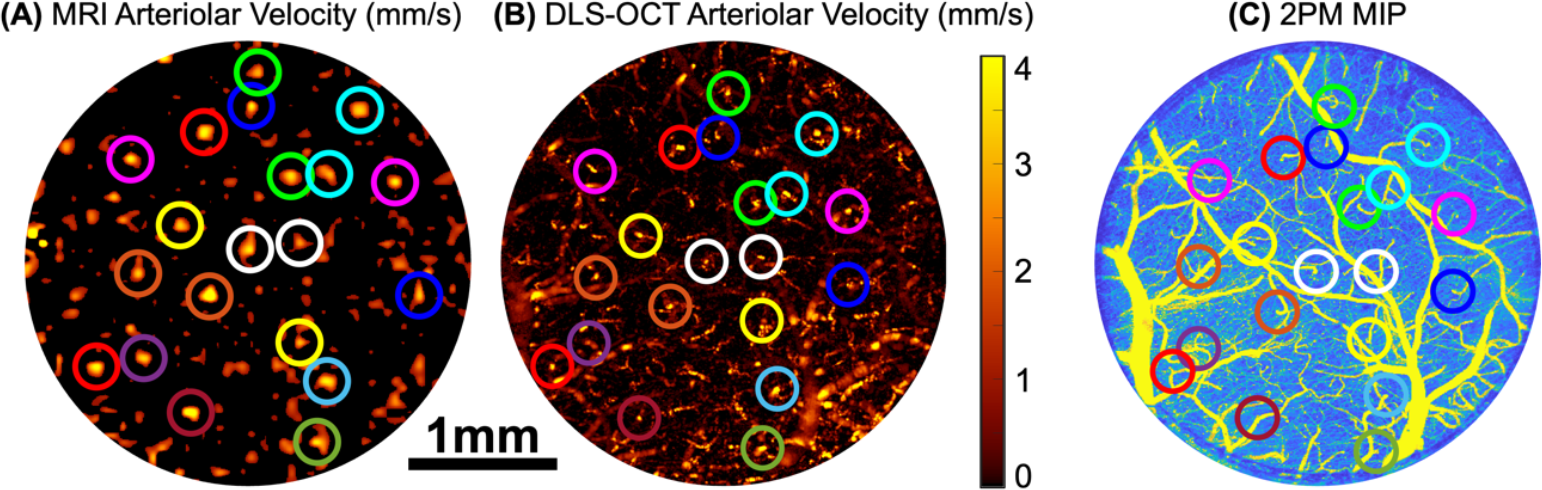
Comparison of arteriolar axial velocity maps from **(A)** PC MRI and **(B)** DLS-OCT for the second mouse. The PC MRI velocity map is a single slice from 50-350 μm below the cortical surface and the DLS-OCT velocity map is a maximum intensity projection (MIP) from 0-500 μm which is why pial vessels are visible. The selected points are also shown in the 2PM microvascular angiogram shown as a MIP from 0-600 μm below the cortical surface **(C)**. The same circle colors in all three panels are used to mark the same diving arterioles. Scale bar is 1 mm and applies to each panel.

## Notes

### Competing Interest Statement

The authors have declared no competing interest.

